# Effective Dynamic Models of Metabolic Networks

**DOI:** 10.1101/047316

**Authors:** Michael Vilkhovoy, Mason Minot, Jeffrey D. Varner

## Abstract

Mathematical models of biochemical networks are useful tools to understand and ultimately predict how cells utilize nutrients to produce valuable products. Hybrid cybernetic models in combination with elementary modes (HCM) is a tool to model cellular metabolism. However, HCM is limited to reduced metabolic networks because of the computational burden of calculating elementary modes. In this study, we developed the hybrid cybernetic modeling with flux balance analysis or HCM-FBA technique which uses flux balance solutions instead of elementary modes to dynamically model metabolism. We show HCM-FBA has comparable performance to HCM for a proof of concept metabolic network and for a reduced anaerobic *E. coli* network. Next, HCM-FBA was applied to a larger metabolic network of aerobic *E. coli* metabolism which was infeasible for HCM (29 FBA modes versus more than 153,000 elementary modes). Global sensitivity analysis further reduced the number of FBA modes required to describe the aerobic *E. coli* data, while maintaining model fit. Thus, HCM-FBA is a promising alternative to HCM for large networks where the generation of elementary modes is infeasible.

## I. INTRODUCTION

Biotechnology harnesses the power of metabolism to produce products that benefit society. Constraints based models are important tools to understand and ultimately to predict how cells utilize nutrients to produce products. Constraints based methods such as flux balance analysis (FBA) [1] and network decomposition approaches such as elementary modes (EMs) [2] or extreme pathways (EPs) [3] model intracellular metabolism using the biochemical stoichiometry and other constraints such as thermodynamical feasibility under pseudo-steady state conditions. FBA has been used to efficiently estimate the performance of metabolic networks of arbitrary complexity, including genome scale networks, using linear programming [4]. On the other hand, EMs (or EPs) catalog all possible metabolic behaviors such that any flux distribution predicted by FBA is a convex combination of the EMs (or EPs) [5]. However, the calculation of EMs (or EPs) is computationally expensive and currently infeasible for genome scale networks [6].

Cybernetic models are an alternative to the constraints based approach which hypothesize that metabolic control is the output of an optimal decision. Cybernetic models have predicted mutant behavior [7, 8], steady-state multiplicity [9], strain specific metabolism [10], and have been used in bioprocess control applications [11]. Hybrid cybernetic models (HCM) have addressed earlier shortcomings of the approach by integrating cybernetic optimality concepts with EMs. HCMs dynamically choose combinations of biochemical modes (each catalyzed by a pseudo enzyme whose expression is controlled by an optimal decision) to achieve a physiological objective (Fig. 1A). HCMs generate intracellular flux distributions consistent with other approaches such as metabolic flux analysis (MFA), and also describe dynamic extracellular measurements superior to dynamic FBA (DFBA) [12]. However, HCMs are restricted to networks which can be decomposed into EMs (or EPs). In this study, we developed the hybrid cybernetic modeling with flux balance analysis (HCM-FBA) technique. HCM-FBA is a modification of the hybrid cybernetic approach of Ramkrishna and coworkers [12] which uses FBA solutions (instead of EMs) in conjunction with cybernetic control variables to dynamically simulate metabolism. Since HCM showed superior performance to DFBA, we compared the performance of HCM-FBA to HCM for a prototypical metabolic network, along with two real-world *E. coli* applications. HCM-FBA performed comparably to HCM for the prototypical network and a reduced anaerobic *E. coli* network, despite having fewer parameters in each case. Next, HCM-FBA was applied to an aerobic *E. coli* metabolic network that was infeasible for HCM. HCM-FBA described cellmass growth and the shift from glucose to acetate consumption with only a few modes. Global sensitivity analysis allowed us to further reduce the aerobic *E. coli* HCM-FBA model to the minimal model required to describe the data. Thus, HCM-FBA is a promising approach for the development of reduced order dynamic metabolic models and a viable alternative to HCM or DFBA, especially for large networks where the generation of EMs is infeasible.

**Fig. 1.**
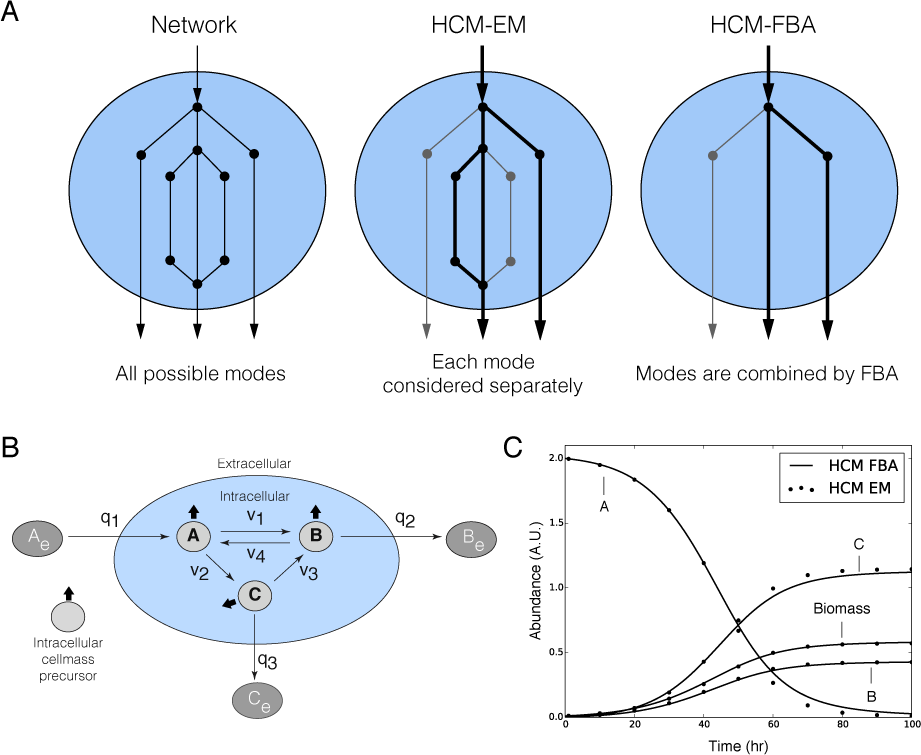
HCM proof of concept metabolic study. A: HCMs distribute uptake and secretion fluxes amongst different pathways. For HCM, these pathways are elementary modes; for HCM-FBA these are flux balance analysis solutions. HCM combines all possible modes within a network; whereas HCM-FBAcombines only steady-state paths estimated by flux balance analysis. B:Prototypical network with six metabolites and seven reactions. Intracellular cellmass precursors *A, B*, and *C* are balanced (no accumulation) while the extracellular metabolites (*A_e_, B_e_*, and *C_e_*) are not balanced (can accumulate). The oval denotes the cell boundary, *q_j_* is the *j*th flux across the boundary, and *v_k_* denotes the *k*th intracellular flux. C: Simulation of extracellular metabolite trajectories using HCM-FBA (solid line) versus HCM (points) for the prototypical network.

## II. RESULTS

HCM-FBA was equivalent to HCM for a prototypical metabolic network (Fig. 1). The proof of concept network, consisting of 6 metabolites and 7 reactions (Fig. 1B), generated 3 FBA modes and 6 EMs. Using the EMs and synthetic parameters, we generated test data from which we estimated the HCM-FBA model parameters. The best fit HCM-FBA model replicated the synthetic data (Fig. 1C). The HCM and HCM-FBA kinetic parameters were not quantitatively identical, but had similar orders of magnitude; the FBA approach had 3 fewer modes, thus identical parameter values were not expected. The HCM-FBA approach replicated synthetic data generated by HCM, despite having 3 fewer modes. Thus, we expect HCM-FBA will perform similarly to HCM, despite having fewer parameters. Next, we tested the ability of HCM-FBA to replicate real-world experimental data.

The performance of HCM-FBA was equivalent to HCM for anaerobic *E. coli* metabolism (Fig. 2A). We constructed an anaerobic *E. coli* network [12], consisting of 12 reactions and 19 metabolites, which generated 7 FBA modes and 9 EMs. HCM reproduced cellmass, glucose, and byproduct trajectories using the kinetic parameters reported by Kim *et al.* [12] (Fig. 2A, points versus dashed). HCM-FBA model parameters were estimated in this study from the Kim *et al.* data set using simulated annealing. Overall, HCM-FBA performed within 5% of HCM (on a residual standard error basis) for the anaerobic *E. coli* data (Fig. 2A, solid), despite having 2 fewer modes and 4 fewer parameters (17 versus 21 parameters). Thus, while both HCM and HCM-FBA described the experimental data, HCM-FBA did so with fewer modes and parameters.

**Fig. 2.**
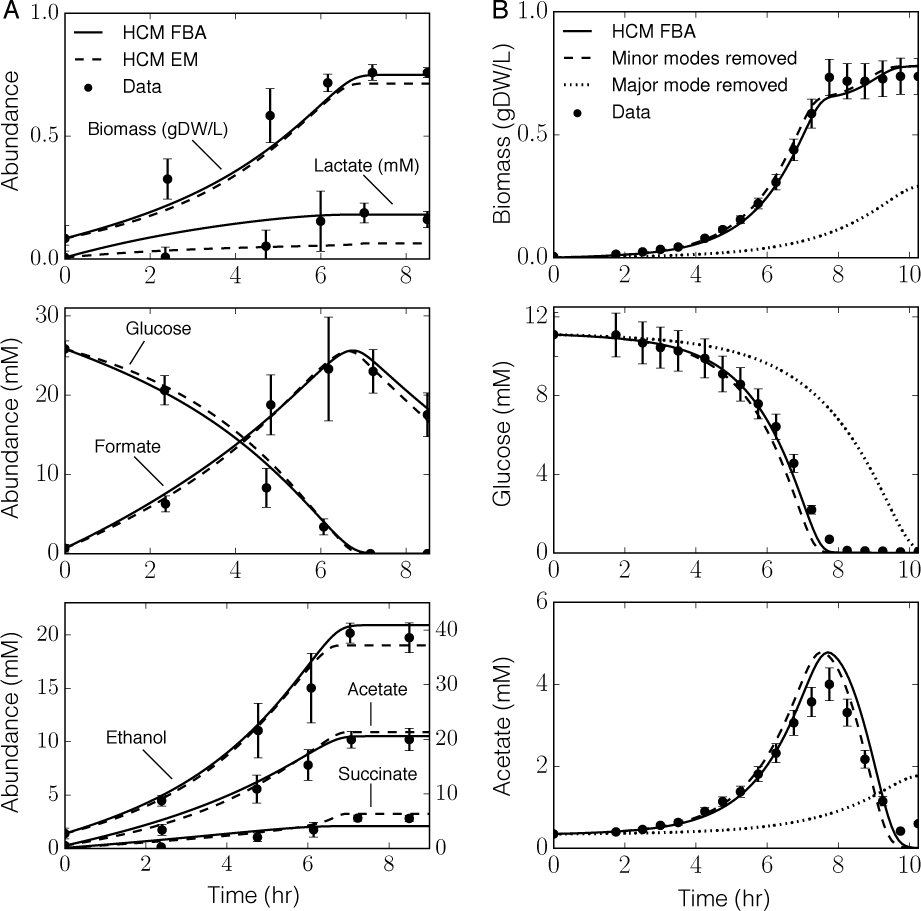
HCM-FBA versus HCM performance for small and large metabolic networks. A: Batch anaerobic *E. coli* fermentation data versus HCM-FBA (solid) and HCM (dashed). The experimental data was reproduced from Kim *et al*. [12]. Error bars represent the 90% confidence interval. B: Batch aerobic *E. coli* fermentation data versus HCM-FBA (solid). Model performance isalso shown when minor modes (dashed) and major modes (dotted) were removed from the HCM-FBA model. The experimental data was reproduced from Varma & Palsson [13]. Error bars denote a 10% coefficient of variation.

HCM-FBA captured the shift from glucose to acetate consumption for a model of aerobic *E. coli* metabolism that was infeasible for HCM (Fig. 2B). An *E. coli* metabolic network (60 metabolites and 105 reactions) was constructed from literature [14,15]. Elementary mode decomposition of this network (and thus HCM) was not feasible; 153,000 elementary modes were generated before the calculation became infeasible. Conversely, flux balance analysis generated only 29 modes for the same network. HCM-FBA model parameters were estimated from cellmass, glucose, and acetate measurements [13] using simulated annealing (Fig. 2B, solid). HCM-FBA captured glucose consumption, cellmass formation, and the switch to acetate consumption following glucose exhaustion. HCM-FBA described the dynamics of a network that was infeasible for HCM, thereby demonstrating the power of the approach for large networks. Next, we demonstrated a systematic strategy to identify the critical subset of FBA modes required for model performance.

Global sensitivity analysis identified the FBA modes essential to model performance (Fig. 3). Total order sensitivity coefficients were calculated for all kinetic parameters and enzyme initial conditions in the aerobic *E. coli* model. Five of the 29 FBA modes were significant; removal of the most significant of these modes (encoding aerobic growth on glucose) destroyed model performance (Fig. 2B, dotted). Conversely, removing the remaining 24 modes simultaneously had a negligible effect upon model performance (Fig. 2B, dashed). The sensitivity analysis identified the minimal model structure required to explain the experimental data.

**Fig. 3.**
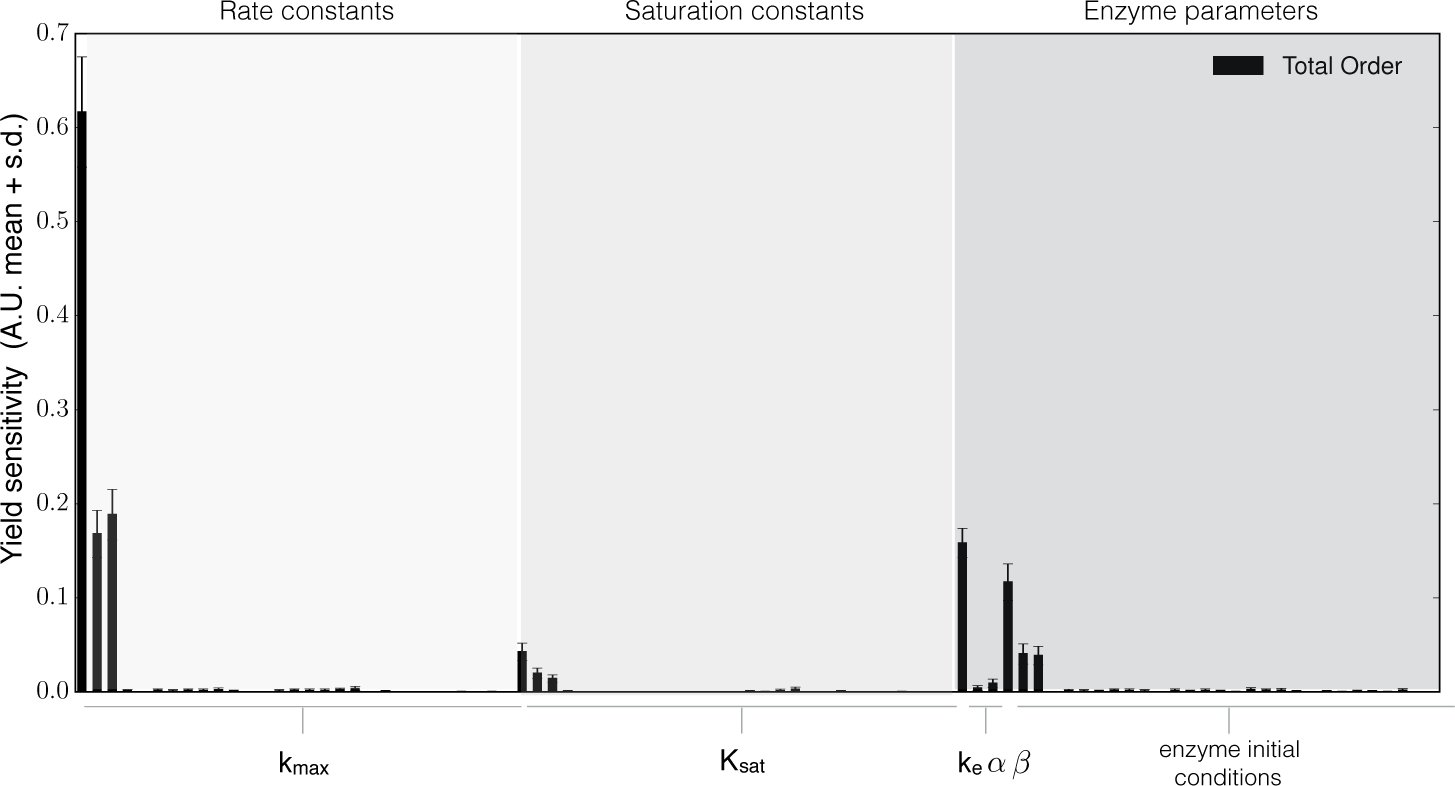
Global sensitivity analysis of the aerobic *E. coli* model. Total order variance based sensitivity coefficients were calculated for the biomass yield on glucose and acetate. Sensitivity coefficients were computed for kinetic parameters and enzyme initial conditions (N = 183,000). Error bars represent the 95% confidence intervals of the sensitivity coefficients.

## III. DISCUSSION

In this study, we developed HCM-FBA, an effective modeling technique to simulate metabolic dynamics. HCM-FBA uses flux balance analysis solutions in conjunction with cybernetic control variables to dynamically simulate metabolism. We studied the performance of HCM-FBA on a prototypical metabolic network, along with two *E. coli* networks. First, we showed that the performance of HCM-FBA and HCM were comparable for the prototypical network and a small model of anaerobic *E. coli* metabolism. For the anaerobic case, both approaches described the experimental data. However, HCM-FBA (which was within 5% of HCM and slightly better than HCM for lactate secretion) had fewer modes and parameters. Next, HCM-FBA was applied to an aerobic *E. coli* metabolic network that was not feasible for HCM. Elementary mode decomposition of the aerobic network generated over 153,000 elementary modes. Conversely, the HCM-FBA approach described cellmass growth and the shift from glucose to acetate consumption with only 29 FBA modes. Global sensitivity analysis further showed that only 5 of the 29 FBA modes were critical to model performance. Removal of these modes crippled the model, but removal of the remaining 24 modes had a negligible impact. These insignificant modes were associated with maintenance, thus they would likely not impact model predictions for a growing culture. HCM-FBA is an alternative approach to HCM, especially for large networks where the generation of elementary modes is infeasible. Elementary modes show the complexity of a cell, displaying the many routes it can take but mathematically FBA has an objective superiority for large networks.

HCM-FBA is a promising approach to model large metabolic networks where elementary modes calculations are infeasible, and where kinetic models of such systems have intractable identification problems. However, there are additional studies that should be performed. First, the intracellular flux distribution predicted by HCM-FBA should be compared to HCM and to flux measurements calculated using MFA or FBA/DFBA in combination with carbon labeling. HCM predicted intracellular fluxes that were similar to MFA results [12]; however, the fluxes predicted by HCM-FBA have not yet been validated. Next, the performance of HCM-FBA should be compared to lumped hybrid cybernetic models (L-HCM). L-HCMs, which combine elementary modes into mode families based upon metabolic function [10, 16], have been applied to an *E. coli* network with 67 reactions and a *Saccharomyces cerevisiae* network with 70 reactions; both cases had satisfactory fits to extracellular experimental data. However, while L-HCM reduces the dimension of possible alternative modes that must be considered, it still requires the calculation of an initial set of modes. For metabolic networks of even moderate size, EM (or EP) decomposition may not be possible. On the other hand, the generation of flux balance solutions (convex combinations of the elementary modes or extreme pathways) is trivial, even for genome scale metabolic networks. Thus, HCM-FBA opens up the possibility for dynamic genome scale models of bacterial and perhaps even of mammalian metabolism.

## IV. MATERIALS AND METHODS

The HCM-FBA approach is a modification of HCM, where elementary modes are replaced with flux balance analysis solutions. Thus, extracellular variables are dynamic while intracellular metabolites are at a pseudo steady state. The abundance of extracellular species *i* (*x_i_*), the pseudo enzyme *e_l_* (catalyzes flux through mode *l*), and cellmass are governed by:

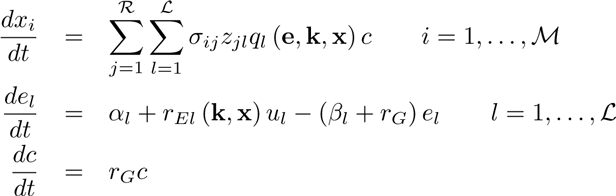

where *ℛ* and *ℳ* denote the number of reactions and extracellular species in the model and *ℒ* denotes the number of FBA modes. The quantity *σ_ij_*, denotes the stoichiometric coefficient for species *i* in reaction *j* and *z_jl_* denotes the normalized flux for reaction *j* in mode *l*. If *σ_ij_* > 0, species *i* is produced by reaction *j*; if *σ_ij_* < 0, species *i* is consumed by reaction *j*; if *σ_ij_* = 0, species *i* is not connected with reaction *j*. Extracellular species balances were subject to the initial conditions x (*t_o_*) = x_*o*_ determined from experimental data. The term *ql* (e, k, x) denotes the specific uptake/secretion rate for mode *l* where e denotes the pseudo enzyme vector, k denotes the unknown kinetic parameter vector, x denotes the extracellular species vector, and *c* denotes the cell mass; *q_l_* (e, k, x) is the product of a kinetic term 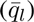 and a control variable governing enzyme activity. Flux through each mode was catalyzed by a pseudo enzyme *e_l_*, synthesized at the regulated specific rate *r_E,l_* (k, x), and constitutively at the rate *α_l_*. The term *u_l_* denotes the cybernetic variable controlling the synthesis of enzyme *l*. The term *β_l_* denotes the rate constant governing non-specific enzyme degradation, and *r_G_* denotes the specific growth rate through all modes. The specific uptake/secretion rates and the specific rate of enzyme synthesis were modeled using saturation kinetics. The specific growth rate was given by:

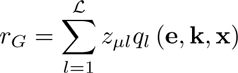

where *z_μl_* denotes the growth flux *μ*, through mode *l*. The control variables *μ_l_* and *v_l_*, which control the synthesis and activity of each enzyme respectively, were given by:

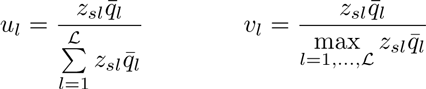

where *z_sl_* denotes the uptake flux of substrate *s* through mode *l*. The model equations were implemented in Julia (v.0.4.2) [17] and solved using SUNDIALS [18]. The model code is available at http://www.varnerlab.org under a MIT license.

*Elementary mode and flux balance analysis:* Elementary modes were calculated using METATOOL 5.1 [19]. FBA modes were defined as the solution flux vector through the network connecting substrate uptake to cellmass and extracellular product formation. The FBA problem was formulated as:

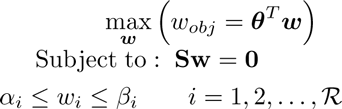

where S denotes the stoichiometric matrix, w denotes the unknown flux vector, *θ* denotes the objective selection vector and *α_i_* and *β_i_* denote the lower and upper bounds on flux *w_i_*, respectively. The flux balance analysis problem was solved using the GNU Linear Programming Kit (v4.52) [20]. For each FBA mode, the objective *w_obj_* was to maximize either the specific growth rate or the specific rate of byproduct formation. Multiple FBA modes were calculated for each objective by allowing the oxygen and nitrate uptake rates to vary. For aerobic metabolism, the specific oxygen and nitrate uptake rates were constrained to allow a maximum flux of 10 mM/gDWhr and 0.05 mM/gDWhr, respectively. Each FBA mode was normalized by the specified objective flux.

*Global sensitivity analysis:* Variance based sensitivity analysis was used to estimate which FBA modes were critical to model performance. The performance function used in this study was the biomass yield on substrate. Candidate parameter sets (N = 182,000) were generated using Sobol sampling by perturbing the best fit parameter set ±50% [21]. Model performance, calculated for each of these parameter sets, was then used to estimate the total-order sensitivity coefficient for each model parameter.

*Estimation of model parameters:* Model parameters were estimated by minimizing the difference between simulations and experimental measurements (squared residual):

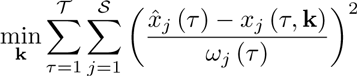

where 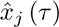 denotes the measured value of species *j* at time *τ*, *x_j_* (*τ*, k) denotes the simulated value for species *j* at time *τ*, and *ω_j_*(*τ*) denotes the experimental measurement variance for species *j* at time *τ*. The outer summation is with respect to time, while the inner summation is with respect to state. The model residual was minimized using simulated annealing implemented in the Julia programming language.

